# Pallidal beta oscillations underlying locomotor adaptation in Parkinson’s disease

**DOI:** 10.64898/2026.08.25.744491

**Authors:** Julia T Choi, Anjela Gurrala, Doris D Wang, Coralie de Hemptinne, Joshua K Wong

## Abstract

**Background:** Locomotor adaptation is essential for adjusting walking patterns to complex environments. This study investigated locomotor adaptation deficits in people with Parkinson’s disease (PD) and examined oscillatory activity in the globus pallidus internus (GPi) during walking adaptation. We hypothesized that elevated beta-band activity in the GPi is associated with reduced locomotor adaptability in PD.

**Methods:** Twelve PD patients with GPi deep brain stimulation (DBS) (eleven bilateral and one unilateral) were included. Local field potentials (LFPs) were recorded from DBS electrodes during split-belt treadmill walking. Patients were tested in the medication-off, DBS-off state. Locomotor adaptation was measured as the change in step length asymmetry during split-belt walking, with smaller changes indicating greater adaptation deficits.

**Results:** We found that GPi high beta (20-30 Hz) and low gamma (30-60 Hz) oscillations were modulated during split-belt walking. Compared to adapters, non-adapters showed decreased movement-related beta suppression during walking. Across participants, beta activity in the GPi contralateral to the fast leg was negatively associated with adaptation magnitude (Spearman’s ρ = −0.65 to −0.75).

**Conclusions:** GPi oscillations are dynamically modulated during locomotor adaptation in PD. Increased beta activity may underlie impaired sensorimotor adaptation during walking. These findings provide novel insight into basal ganglia mechanisms of gait adaptation in PD and suggest that elevated GPi beta activity may serve as a marker of locomotor adaptation deficits.

## Introduction

Locomotor adaptation is a fundamental form of sensorimotor learning that enables adjustments in walking patterns to changing environmental demands.^1^ The split-belt treadmill paradigm measures locomotor adaptation from changes in gait asymmetry during adaptation and after-effects during post-adaptation. ^2^ In healthy adults, step length asymmetry initially becomes negative and gradually returns to baseline.^2^ Previous studies using the split-belt paradigm have demonstrated substantial individual variability in locomotor adaptation across people with Parkinson’s disease (PD), with some individuals showing little to no adaptation (“non-adapters”).^3–5^ Impaired locomotor adaptation in PD has been linked to cognitive deficits and freezing of gait (FOG).^5–8^ Although individuals with FOG often exhibit greater deficits in locomotor adaptation, impaired adaptation is not exclusive to freezers.^3^ Further research is needed to understand the pathophysiology underlying locomotor adaptation deficits in PD.

The basal ganglia-thalamocortical loops are essential for higher-order regulation of gait control, while the basal ganglia-brainstem projections influence muscle tone and posture, together ensuring adaptive and stable walking.^9^ The globus pallidus internus (GPi) is a major basal ganglia output center and a target for deep brain stimulation (DBS) in Parkinson’s disease.^10–12^ Anatomically, the GPi is centrally positioned to modulate gait given its direct connections to both the thalamocortical network and brainstem locomotor regions. In Parkinson’s disease, overactive inhibitory output from the GPi is hypothesized to play a role in delayed gait initiation, reduced gait velocity, and the manifestation of FOG.^13^

Communication between cortical and subcortical networks is supported by neural oscillations at different frequencies, reflecting synchronized activity within and between neuronal networks.^14^ Local field potentials (LFPs) recorded via DBS electrodes have demonstrated that the basal ganglia exhibit gait-related oscillatory dynamics, including suppression of beta oscillations, shorter beta bursts, and reduced bilateral connectivity during walking.^15–17^ During overground walking, subthalamic nucleus oscillations are time-locked to leg muscle activation patterns, and exhibit gait phase-dependent coherence with the sensorimotor cortex.^18, 19^ Excessive synchronization in the basal ganglia, particularly in the beta frequency (13-35 Hz) range, has been suggested to serve as a marker for distinguishing normal vs. pathological gait in PD.

Understanding these basal ganglia oscillations and how they change under different walking conditions is important for therapeutic interventions.

We studied pallidal oscillatory activity during locomotor adaptation in people with PD using GPi LFPs recorded from bidirectional DBS electrodes during split-belt walking. Patients were tested in the medication-off, DBS-off state. Adaptation was quantified by changes in step length asymmetry, a measure reflecting spatial (foot placement) and temporal (step time) control strategies.^20^ We hypothesized that 1) GPi beta activity would be modulated walking adaptation, and that 2) elevated beta oscillations would be associated with impaired locomotor adaptation.

## Materials and methods

### Participants

Patients diagnosed with idiopathic Parkinson’s disease based upon the Movement Disorders Society clinical diagnostic criteria,^21^ who underwent bilateral (n=11) or unilateral (n=1) GPi-DBS were included in this study (**Table 1**). All patients were implanted with a Medtronic Percept PC neurostimulator and either Medtronic 3387 or SenSight B33015 DBS leads (Medtronic, Minneapolis, MN). This study included patients with chronic, stable DBS defined as having completed at least six months of DBS optimization at the Normal Fixel Institute for Neurological Diseases and maintaining stable DBS settings for at least one month. FOG status was determined by any clinical documentation of freezing episodes, whether observed during clinic visits or reported to occur at home. All participants provided written informed consent prior to the study, in accordance with the Declaration of Helsinki and the protocol approved by the Institutional Review Board of the University of Florida, Gainesville, FL (IRB # 20192465).

**Table 1.** Demographics.

| <sup>a</sup>Subject | Age | Sex | GPI-DBS | Hoehn and Yahr Stage | <sup>b</sup>UPDRS III | <sup>c</sup>FOG status | <sup>d</sup>Adapter status | <sup>e</sup>Fast gait speed, m/s |
| --- | --- | --- | --- | --- | --- | --- | --- | --- |
| 001 | 71 | F | bilateral | 3 | 24 | Freezer | Non-adapter | 0.8 (right) |
| 002 | 60 | F | bilateral | 2.5 | 26 | Freezer | Adapter | 0.8 (right) |
| 003 | 59 | F | bilateral | 3 | 33 | Freezer | Non-adapter | 0.2 (left) |
| 004 | 67 | M | left | 2 | 24 | Freezer* | Non-adapter | 0.4 (right) |
| 005 | 40 | M | Bilateral<sup>§</sup> | 2.5 | 24 | Non-freezer | Adapter | 0.6 (right) |
| 007 | 69 | M | bilateral | 3 | 56 | Freezer | Non-adapter | 0.6 (right) |
| 008 | 72 | M | bilateral | 2 | 15 | Freezer | Adapter | 0.8 (right) |
| 009 | 63 | M | bilateral | 2 | 12 | Non-freezer | Adapter | 1.0 (left) |
| 011 | 67 | M | bilateral | 2.5 | 29 | Non-freezer | Adapter | 0.6 (right) |
| 012 | 45 | M | bilateral | 2 | 21 | Non-freezer | Adapter | 1.2 (right) |
| 013 | 67 | F | bilateral | 3 | 29 | Freezer | Adapter | 0.9 (left) |
| 014 | 53 | F | Bilateral<sup>§</sup> | 2.5 | 14 | Non-freezer | Non-adapter | 0.6 (left) |

a Subject IDs ranged from 001 to 014. This analysis only included those with GPI-DBS. Two participants with STN-DBS were not included.
b MED ON DBS ON scores within +/- 12 months of the study visit, except for 0011 which is 14 months from the study visit.
c Patients were classified as freezers or non-freezers based on their clinical history. \* Subject 004 exhibited FOG during the experimental session.
d Adapters and non-adapters were classified based on participant's change in step length asymmetry during split-belt treadmill walking
e Participant's fastest comfortable treadmill speed. The leg assigned to the fast belt during split-belt adaptation is indicated in parentheses.
<sup>§</sup> Right LFPs from Subject 005 and Left LFPs from Subject 014 were excluded from analysis due to electrocardiographic artifacts

### Experimental Protocol

Patients were tested after overnight withdrawal of dopaminergic medication. DBS remained activated overnight and was discontinued in the laboratory to obtain a clinical off-state (medication OFF, stimulation off), with a 20-minute DBS washout period before testing.

A rest recording was conducted, during which they sat comfortably and relaxed with eyes open. Participants then walked on an instrumented split-belt treadmill (Bertec Corporation, Columbus, OH, USA), wearing a ceiling-mounted safety harness (Solo-Step Inc., North Sioux City, SD, USA). The range of walking speed was first explored by starting the treadmill at 0.2 m/s and gradually increasing it until the subject reported reaching their fastest comfortable walking speed. This was defined as the ‘fast speed’, while the ‘slow speed’ was set to half of the fast speed.

The split-belt walking consisted of three phases (**Fig. 1a**). 1) During baseline, participants walked at the slow speed for 2 minutes. 2) This was followed by a 6-minute adaptation, where one belt moved at the fast speed and the other at the slow speed, creating a speed asymmetry. The assignment of fast leg was randomized across participants. 3) During post-adaptation, the belts moved at the slow speed for 3 minutes. The treadmill was stopped briefly before changing conditions.

**Figure 1.**
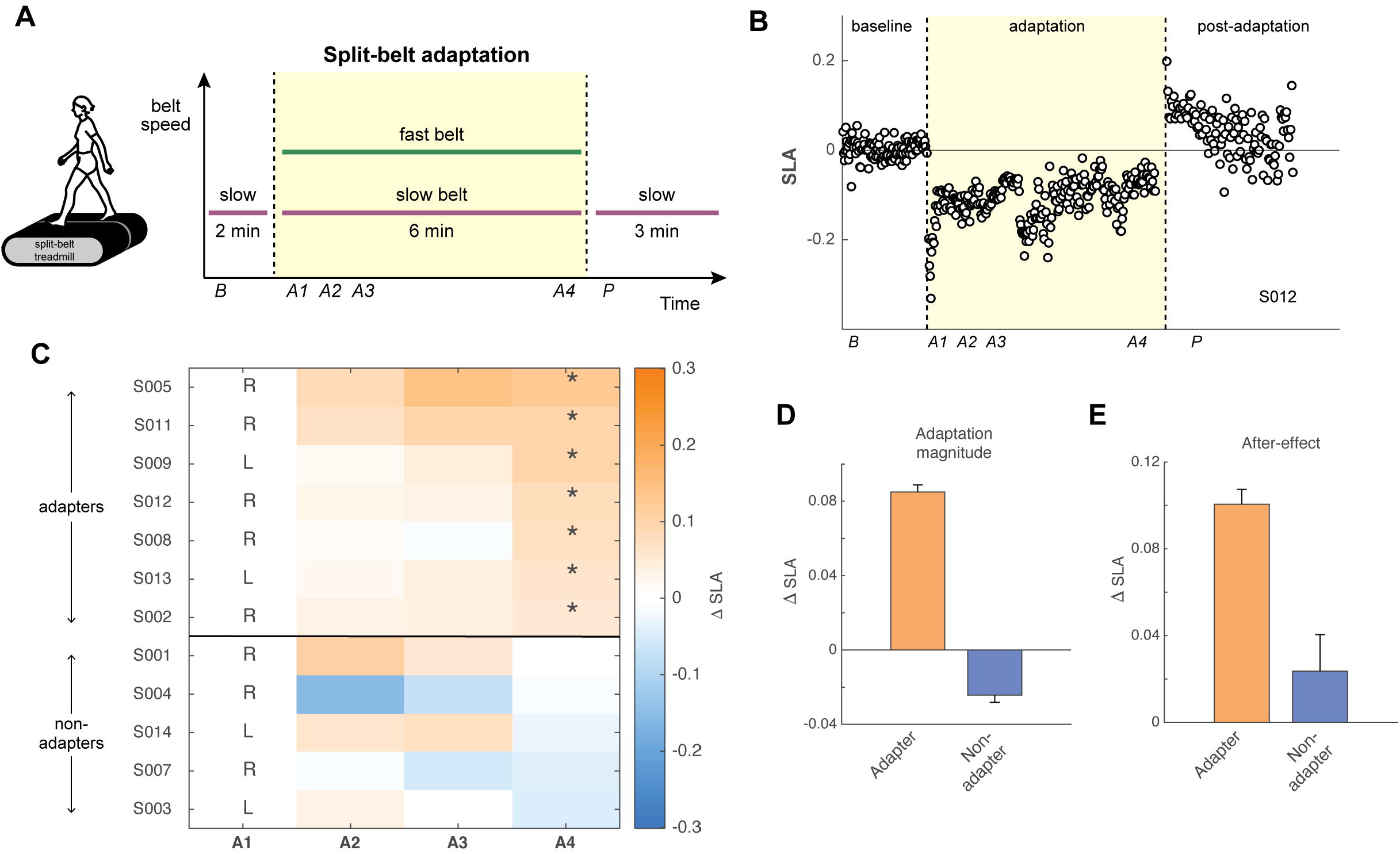
Step length adaptation during split-belt walking. **(A)** Experimental paradigm illustrating treadmill belt speeds across baseline, adaptation and post-adaptation phases. Analysis epochs were defined as baseline (*B*); early adaptation *A1* (0-30 s), *A2* (30-60 s), *A3* (60-90 s), and late adaptation *A4* (last 30 s of adaptation); and post-adaptation (*P*). **(B)** Stride by stride step length asymmetry (SLA) over time for a representative “adapter”. Zero indicates symmetric step lengths. **(C)** Change in SLA for each participant across adaptation epochs. ΔSLA is expressed relative to *A1*. More positive values indicate greater adaptation. Rows are sorted in descending order of adaptation magnitude. R and L denote the leg assigned to the fast belt during adaptation. Asterisks (^*^) indicates a significant difference between *A4* and *A1*. Horizontal black line separates adapters from non-adapters. **(D)** Group differences in adaptation magnitude. **(E)** Group differences in size of after-effects.

### Data collection

Reflective markers were placed bilaterally on the toe (fifth metatarsal), ankle, knee, hip and anterior superior iliac spine. Marker data were recorded at 100 Hz using Miqus cameras (Qualisys AB, Gothenburg, Sweden). Ground reaction forces (GRF) were sampled at 1000 Hz. A sync unit was used to trigger simultaneous start of kinematics and force recordings.

LFPs were recorded at 250 Hz using the BrainSense Indefinite Streaming mode (Medtronic Inc., Minneapolis, MN, USA), for three bipolar contact pairs (C0-C3, C1-C3, C0-C2) per hemisphere. Prior to recording, the internal clock of the implanted pulse generator was synchronized to match that of the Qualisys system.

### Data processing

Signal processing was performed in MATLAB (R2023b, MathWorks Inc., Natick, MA, USA). GRFs were low-pass filtered at 15 Hz using a zero-phase 3^rd^ order Butterworth filter. Stance was defined by vertical GRF > 2% body weight (i.e., foot contact to toe-off). Gait events were visually inspected and corrected as needed.

Step length was calculated as the anterior-posterior distance between ankle markers at foot contact. Step length asymmetry (SLA) was calculated as the normalized different in step length between two legs:

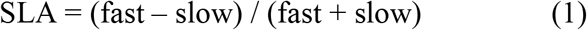

where a value of zero indicates perfect symmetry, positive values indicate longer steps with the fast leg, and negative values indicate longer steps with the slow leg.

Adaptation magnitude was quantified as the change in SLA from the first 30 to the last 30 seconds of the split-belt adaptation period. After-effects were assessed as the difference in SLA during the first 30 seconds of post-adaptation compared to baseline.

LFP analyses were based on the C0–C3 contact pair, as this widest bipolar configuration maximizes spatial coverage across the GPi while avoiding signal redundancy from the overlapping C1–C3 and C0–C2 pairs. Due to cardiac artifacts, only left hemisphere from Subject 5 and right hemisphere from Subject 14 were analyzed. Raw LFP recordings were band-pass filtered (1-100 Hz). Power spectral density (PSD) was computed using Welch’s method (0.5 Hz resolution, 50% overlap). Power was averaged over the theta (4-8 Hz), alpha (8-13 Hz), low beta (13-20 Hz), high beta (20-30 Hz) and low gamma (30-60 Hz) bands. Spectral power values were normalized by subtracting the resting-state power, with reported values representing the change relative to rest.

### Statistical analysis

Changes in SLA and LFP power were analyzed across the following epochs: baseline (*B*: first 30 s of slow walking), adaptation (*A1*: 0-30 s, *A2*: 30-60 s, *A3*: 60-90 s, and *A4*: final 30 s), and post-adaptation (*P*: first 30 s). Adaptation epochs were selected to capture both early adaptation dynamics and late steady-state behavior.^22^

A repeated-measures ANOVA was used to determine changes in SLA, with epoch as within-subject factor. Because the assumption of sphericity was not met for epoch (Mauchly’s χ^2^ =32.378, *p* = 0.005); degrees of freedom were corrected using Greenhouse-Geisser. Bonferroni-corrected post hoc comparisons were conducted to identify differences between epochs.

For each participant, a one-tailed t-test was used to determine whether SLA became less negative during adaptation (*A4* compared to *A1)*. Participants were classified as “adapters” if the *p*-value was less than 0.05. A separate one-tailed t-test was used to determine whether post-adaptation SLA was more positive than baseline, indicating the presence of after-effects. Group differences were assessed using Welch’s t-tests for continuous variables and Fisher’s exact test for categorical variables.

Linear mixed-effects models were implemented in R (*lme4, lmerTest*) to examine the effects of Side (ipsilateral vs. contralateral relative to the fast leg), Epoch, and their interaction on LFP power within each frequency band. Adaptation Status (adapter vs. non-adapter) and FOG status (freezer vs. non-freezer) were included as additional fixed main effects. Models included subject-specific random intercepts and random slopes for Side. Fixed effects were evaluated using Type III ANOVA with Satterthwaite’s approximation for degrees of freedom. Post hoc pairwise comparisons were performed using estimated marginal means (*emmeans*) with Kenward–Roger correction and Tukey adjustment for multiple comparisons.

Spearman correlations were used to examine the relationship between LFP power and adaptation magnitude. Because walking speed differed across participants, partial correlation analyses controlling for speed were performed to determine whether these relationships were independent of speed.

## Results

### Step length adaptation

There was a significant effect of epoch on SLA (F(0.25,2.42) = 12.00, *p* < 0.001). Relative to baseline, SLA became more negative during early adaptation (*A1*) (Cohens’ *d* = 0.59, *p* =0.03), reflecting shorter steps on the fast leg. Over the course of adaptation, SLA gradually returned toward baseline values and was not different from baseline by late adaptation (*A4)* (*d* = 0.36, *p* = 0.4). This pattern is consistent with normal adaptation during split-belt walking. During post-adaptation, SLA shifted in the positive direction (i.e., after-effects), although this was not significantly different from baseline at the group level (*d* = −0.41, *p* = 0.16).

At the individual level, seven participants showed a significant change in SLA from *A1* to *A4* and were classified as “adapters” (**Fig. 1c**). After-effects were confirmed in all adapters. Five participants did not show a significant change in SLA and were classified as “non-adapters”. The Supplementary Videos show representative walking patterns of an adapter and a non-adapter during split-belt walking.

Baseline SLA was not significantly different between adapter and non-adapter groups (*p* = 0.4). The treadmill speed was higher for adapters (*p* = 0.03), but the number of steps taken during the adaptation did not significantly differ between groups (*p* = 0.8). Although non-adapters were predominantly freezers, the proportion of freezers did not differ significantly between groups (**Table 2**).

**Table 2.** Characteristics by participant groups.

|  | **Adapters** | **Non-adapters** | **p-value** |
| --- | --- | --- | --- |
| Age, yrs | 59 ± 12 | 64 ± 8 | p = 0.4<sup>1</sup> |
| Sex | 5 males: 2 females | 2 males: 3 females | p = 0.6<sup>1</sup> |
| Disease duration, yrs | 11 ± 4 | 15 ± 7 | p = 0.3<sup>1</sup> |
| UPDRS III | 22 ± 7 | 30 ± 16 | p = 0.6<sup>1</sup> |
| FOG status | 3 freezers: 4 non-freezers | 4 freezers: 1 non-freezer | p = 0.3<sup>2</sup> |
| Fast speed, m/s | 0.84 ± 0.21 | 0.52 ± 0.23 | p = 0.03<sup>1</sup> |
| Fast leg | 5 right: 2 left | 3 right: 2 left | p = 1<sup>2</sup> |
| Baseline SLA | -0.012 ± 0.03 | -0.145 ± 0.27 | p = 0.3<sup>1</sup> |
| <sup>a</sup>No. of steps | 448 ± 88 | 441 ± 216 | p = 0.8<sup>1</sup> |

<sup>1</sup> Welch's t-test
<sup>2</sup> Fisher's exact test
SLA = step length asymmetry
<sup>a</sup> No. of steps = number of steps taken during the split-belt adaptation phase

Adaptation magnitude differed significantly between groups (**Fig. 1d**; t(9.97) < −8.17, *p* < 0.001, *d* = −4.66). After-effects were not significantly different between groups, although they tend to be larger in adapters (**Fig. 1e**; t(5.87) < −1.84, *p* = 0.1, *d* = −1.12).

### GPi oscillations during split-belt adaptation

GPi activity was modulated during walking relative to rest (**Fig. 2a,b**). Non-adapters showed greater low beta (F(1,9.84) = 6.46, *p* = 0.03) and high beta power (F(1,7.57) = 14.99, *p* = 0.005), than adapters during walking (**Fig. 2c**). FOG was independently associated with elevated low beta power (F(1,9.84) = 6.73, *p* = 0.027) but not high beta power.

**Figure 2.**
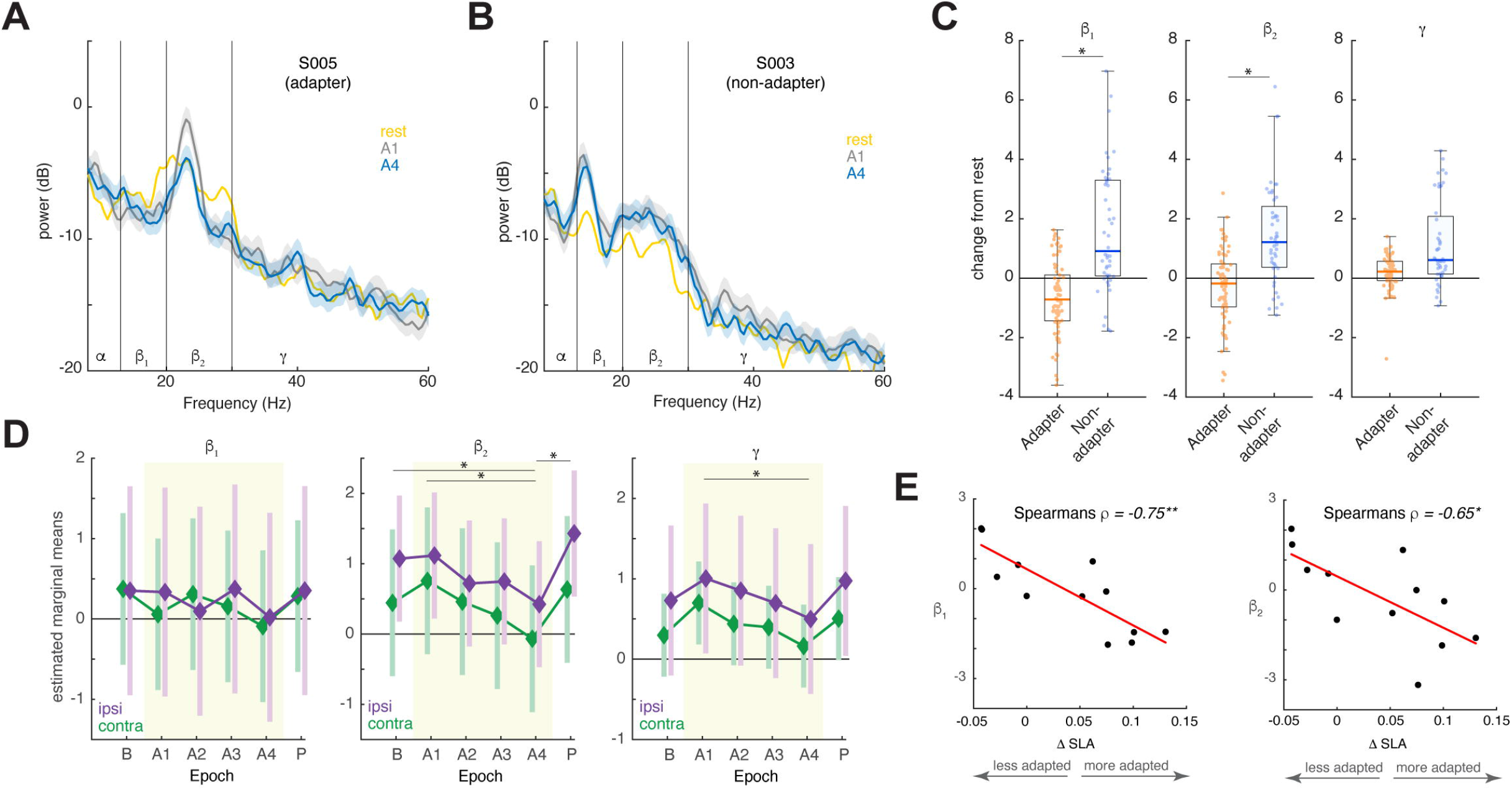
GPi activity during split-belt walking. **(A-B)** Power spectral density (PSD, mean ± 95% confidence interval) of contralateral GPi for an adapter (S005; A) and a non-adaptor (S003; B). **(C)** GPi LFP power relative to rest. Box plots show the aggregated distribution of low beta (β_1_, 13-20 Hz), high beta (β_2_, 20-30 Hz) and low gamma (γ, 30-60 Hz) power in adapters vs. non-adapters. Dots represent values from each participant, epoch, and hemisphere. **(D)** GPi LFP power across walking epochs. Diamond symbols represent estimated marginal means from the linear mixed-effects model, and error bars represent 95% confidence intervals for each hemisphere (contralateral: green, ipsilateral: purple). Adaptation epochs (*A1-A4*) are shaded. Asterisks denote significant difference between epochs (Tukey-adjusted *p* < 0.05). See Supplementary Figure S1 for individual data points across epochs. **(E)** Relationship between ΔSLA and contralateral beta power during late adaptation (*A4*). Spearman’s ρ is shown; *p* < 0.05 (*), *p* < 0.001 (**).

High beta and low gamma power were modulated across epochs (**Fig. 2d**). Linear mixed-effects models revealed a significant effect of epoch for high beta (F(5,95) = 4.90, *p* < 0.001) and low gamma power (F(5,95) = 2.86, *p* = 0.02), but not theta, alpha and low beta bands (all *p* > 0.5). There were no effects of side, or side × epoch interactions (**Table 3**). High beta power decreased at *A4* relative to baseline (*p* < 0.05) and *A1* (*p* = 0.003), followed by a rebound during post-adaptation (*p* < 0.001). Low gamma power was lower at *A4* relative to *A1* (*p* = 0.012), with no other differences between epochs.

**Table 3.** Main effects on LFP power from linear mixed-effect models across frequency bands.

| Fixed Effects | Theta<br>4-7 Hz | Alpha<br>8-13 Hz | Low Beta<br>13-20 Hz | High Beta<br>20-30 Hz | Low Gamma<br>30-60 Hz |
| --- | --- | --- | --- | --- | --- |
| Side | $F(1,8.12) = 1.710$<br>$p = 0.227$ | $F(1,8.58) = 0.090$<br>$p = 0.771$ | $F(1,10.25) = 0.012$<br>$p = 0.914$ | $F(1,11.53) = 0.569$<br>$p = 0.466$ | $F(1,8.63) = 0.830$<br>$p = 0.387$ |
| Epoch | $F(5,95) = 0.379$<br>$p = 0.862$ | $F(5,95) = 0.782$<br>$p = 0.565$ | $F(5,95) = 0.755$<br>$p = 0.584$ | $F(5,95) = 4.898$<br>**$p < 0.001$** | $F(5,95) = 2.860$<br>**$p = 0.019$** |
| Side x Epoch | $F(5,95) = 0.442$<br>$p = 0.818$ | $F(5,95) = 0.155$<br>$p = 0.978$ | $F(5,95) = 0.296$<br>$p = 0.914$ | $F(5,95) = 0.459$<br>$p = 0.806$ | $F(5,95) = 0.111$<br>$p = 0.990$ |
| Adapter status | $F(1,8.99) = 0.611$<br>$p = 0.454$ | $F(1,9.49) = 1.469$<br>$p = 0.255$ | $F(1,9.84) = 6.462$<br>**$p = 0.030$** | $F(1,7.57) = 14.993$<br>**$p = 0.005$** | $F(1,9.30) = 2.777$<br>$p = 0.129$ |
| FOG status | $F(1,8.99) = 1.003$<br>$p = 0.343$ | $F(1,9.49) = 1.144$<br>$p = 0.311$ | $F(1,9.84) = 6.733$<br>**$p = 0.027$** | $F(1,7.57) = 0.114$<br>$p = 0.745$ | $F(1,9.30) = 2.135$<br>$p = 0.177$ |

Contralateral GPi beta power was negatively correlated with adaptation magnitude (**Fig. 2e)**, for both low beta (Spearman’s ρ = −0.75, *p* = 0.008) and high beta frequencies (ρ = −0.65, *p* = 0.03), after controlling for walking speed. Low-gamma power was also negatively correlated with adaptation magnitude (ρ = −0.64, *p* = 0.03); but its independent contribution could not be determined because it also correlated with beta power (ρ = 0.77-0.86, *p* < 0.001). Ipsilateral GPi power was not correlated with adaptation magnitude (all *p* > 0.4).

## Discussion

This study provides novel evidence that oscillatory activity in the GPi is dynamically changing during locomotor adaptation in PD. High beta power was modulated across adaptation epochs and differed by adaptation status, suggesting it reflects neural processes underlying locomotor adaptation. Elevated GPi beta power along with decreased movement related beta suppression was observed in non-adapters, indicating that increased beta activity may underly impaired sensorimotor recalibration during walking in PD.

Pallidal activity during split-belt adaptation was modulated in the high beta and low gamma frequency bands. In healthy individuals, beta-gamma EMG coherence has been shown to increase at the onset of split-belt walking and decreases with adaptation, reflecting modulation of common drive to leg muscles in response to the novel walking environment.^23, 24^ Moreover, beta power in the sensorimotor and parietal cortices has been shown to decrease during the early phase of split-belt walking adaptation and return to baseline by late adaptation.^25^ Together, these findings suggest that modulation of beta-gamma oscillations across cortical and subcortical neural oscillations may be functionally linked during locomotor adaptation.

Contralateral GPi beta activity was significantly correlated with adaptation magnitude, indicating that elevated beta activity is a potential marker of impaired locomotor adaptability. This aligns with the broader framework of movement-related beta suppression, in which beta activity is reduced during normal walking. Prior work has shown beta desynchronization in the subthalamic nucleus (STN) during stepping in place and overground walking in PD.^16, 17^ Non-freezers typically show a reduction in STN beta power across the low-and high-beta frequency bands during walking compared to rest, while freezers show increases in low beta activity during gait.^17, 26^ Relative to STN, GPi dynamics during gait is less understood. Our current findings suggest that elevated GPi beta activity may reflect a common deficit in beta suppression, contributing to impaired gait control and reduced adaptability in PD.^27^

Decreased movement-related beta desynchronization in non-adapters may reflect altered neural circuitry governing error-based motor learning in PD. Adapting to the asymmetric constraints of a split-belt treadmill relies on the continuous resolution of sensory prediction errors, a process classically localized to cerebellar networks.^28^ However, sensorimotor adaptation involves interactions between cerebellar and basal ganglia circuits, with the relative contributions of these networks differing according to task demands and level of expertise.^29^ The lack of GPi beta desynchronization in non-adapters may suggest reduced engagement of the basal ganglia during locomotor adaptation. This is consistent with studies demonstrating increased reliance on cerebellar pathways during motor tasks in PD.^30^ Together, these findings suggest that adaptation deficits may reflect impaired cerebellar-basal ganglia interactions that support adaptive locomotion in PD.

Furthermore, cognitive function may influence the ability to adapt to the split-belt paradigm. Movement-related beta desynchronization is traditionally a hallmark of successful motor planning and execution.^31^ Initial adaptation to the mechanical perturbation of a split-belt treadmill is cognitively demanding, requiring increased attention and ongoing gait adjustments to maintain stability.^32^ Given that beta desynchronization strongly correlates with the degree of executive demand during motor processing,^33^ greater beta suppression would be expected during split-belt walking then unconstrained walking conditions. However, non-adapters did not exhibit beta desynchronization despite the increased task demands of split-belt walking. This may reflect deficits in high-level cognitive processing, where executive dysfunction prevents the basal ganglia from effectively engaging in task-dependent motor planning.

A limitation of this study is that each participant completed only one split-belt configuration, and the belt assignment may have influenced our results. Second, the 20-minute DBS washout period may be insufficient to achieve a complete OFF-DBS state, as prior work suggests that washout of axial symptoms can require several hours,^34^ but comparable data for GPi DBS are unavailable. Finally, split-belt walking differs from overground walking, which may alter gait dynamics and reduce freezing episodes. Although impaired adaptation is more common in individuals with FOG, not all freezers showed deficits, suggesting that reduced adaptability reflects a distinct clinical feature rather than a consequence of FOG. Larger samples are needed to disentangle locomotor adaptation deficits from FOG and identify their neural correlates.

These findings have important implications for neuromodulation strategies targeting gait in PD. Investigating the effects of DBS on locomotor adaptation, particularly in individuals with poor adaptation capacity, will be an important next step. If beta activity in the GPi reflects sensorimotor recalibration, then modulation of this signal may enhance gait adaptability in PD. Recent advances in closed-loop DBS that adjust stimulation based on beta burst activity, activity-dependent adaptive simulation, as well as adaptive stimulation synchronized to specific phases of the gait cycle, highlight the potential for temporally and spectrally precise neuromodulation to improve gait in PD.^35–37^

## Video legends

**Supplementary Video 1**

Animated visualization of split-belt walking during the late adaptation phase in a representative adapter.

**Supplementary Video 2**

Animated visualization of split-belt walking during the late adaptation phase in a representative non-adapter.

## Notes

### Competing Interest Statement

The authors have declared no competing interest.

### Funding

- National Institute on Aging: R21 AG088600
- UF Research Opportunity Seed Fund
- Mangurian-Fixel-McKnight Collaboration Pilot Studies

