## Supplementary Figure S1 for "Pallidal beta oscillations underlying locomotor adaptation in Parkinson’s disease"

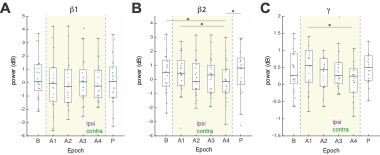

**Supplementary Figure S1. Observed changes in GPi oscillatory power across walking epochs.** Box plots show change in low-beta (β_1_, 13-20 Hz, **A**), high beta (β_2_, 20-30 Hz, **B**) and low-gamma (γ, 30-60 Hz, **C**) power. Colored dots represent individual power values by hemisphere (contralateral: green, ipsilateral: purple). Epochs include baseline (*B*), adaptation (*A1* to *A4*; shaded region), and post-adaptation (*P*). Asterisks denote significant post hoc comparisons (*p* < 0.05).
